# Loss of the conserved Alveolate kinase MAPK2 decouples *Toxoplasma* cell growth from the cell cycle

**DOI:** 10.1101/2020.05.13.095091

**Authors:** Xiaoyu Hu, William J. O’Shaughnessy, Tsebaot G. Beraki, Michael L. Reese

## Abstract

Mitogen-activated protein kinases (MAPKs) are a conserved family of protein kinases that regulate signal transduction, proliferation, and development throughout eukaryotes. The Apicomplexan parasite *Toxoplasma gondii* expresses three MAPKs. Two of these, ERK7 and MAPKL1, have been respectively implicated in the regulation of conoid biogenesis and centrosome duplication. The third kinase, MAPK2, is specific to and conserved throughout Alveolata, though its function is unknown. We used the auxin-inducible degron system to determine phenotypes associated with MAPK2 loss-of-function in *Toxoplasma*. We observed that parasites lacking MAPK2 failed to duplicate their centrosomes and therefore did not initiate daughter-cell budding, which ultimately led to parasite death. MAPKL2-deficient parasites initiated, but did not complete DNA replication, and arrested prior to mitosis. Surprisingly, the parasites continued to grow in size and to replicate their Golgi, mitochondria, and apicoplasts. We found that the failure in centrosome duplication is distinct from the phenotype caused by depletion of MAPKL1. As we did not observe MAPK2 localization at the centrosome at any point in the cell cycle, our data suggest MAPK2 regulates a process at a distal site that is required for completion of centrosome duplication and initiation of parasite mitosis.

**Importance:** *Toxoplasma gondii* is a ubiquitous intracellular protozoan parasite that can cause severe and fatal disease in immunocompromised patients and the developing fetus. Rapid parasite replication is critical for establishing a productive infection. Here, we demonstrate that a *Toxoplasma* protein kinase called MAPK2 is conserved throughout Alveolata and essential for parasite replication. We found that parasites lacking MAPK2 protein were defective in the initiation of daughter cell budding and were rendered inviable. Specifically, TgMAPK2 appears to be required for centrosome replication at the basal end of the nucleus, and its loss causes arrest early in parasite division. MAPK2 is unique to Alveolata and not found in metazoa, and likely is a critical component of an essential parasite-specific signaling network.

## Introduction

Cellular replication depends upon the faithful duplication and partition of genetic material and cell organelles. Successful progression through the eukaryotic cell cycle is controlled by a network of well-conserved regulatory components that not only organize different stages, but also adapt to extracellular signals (1). The obligate intracellular apicomplexan parasites have among the most diverse replicative paradigms among eukaryotes (2). These unusual cell cycles appear to be controlled by mechanisms both analogous to and divergent from their metazoan hosts (3–5). The *Toxoplasma gondii* asexual cycle replicates via “endodyogeny,” wherein two daughters grow within an intact mother (6). Efficient replication of these parasites is critical to establishment of infection and also results in host tissue damage, thus contributing considerably to pathogenesis.

As the physical anchor for DNA segregation, the centrosome is central to the organization of cell division. Centrosome duplication is one of the earliest and most decisive events in the parasite cell cycle (3, 7–10). Earlier work identified two distinct centrosomal cores: an outer core and an inner core which orchestrate cytokinesis and nuclear division independently (3, 11). Daughter cell budding initiates prior to completion of mitosis, and the parasite centrosome is a central hub that coordinates both mitosis and daughter budding. Budding begins at the end of S phase with daughter cytoskeletal components assembling near the duplicated centrosomes, which provide a scaffold for daughter cell assembly (8, 12, 13). Organelles are then partitioned between these elongating scaffolds (14). A growing number of the regulatory components of the parasite cell cycle have been identified, including homologs of scaffolds (4, 15) and kinases (5, 8, 16, 17) that are broadly conserved in other organisms. However, the precise roles these factors play in *Toxoplasma* are often distinct from what they are known to play in model organisms – this is likely due to the parasite’s specialized cell cycle (6) and unusual centrosomal structure (3, 11). Thus, even the study of well-conserved proteins can yield surprising insight into the functional adaptations evolved in the network that regulates the parasite cell cycle.

For example, throughout eukaryotes, members of the mitogen-activated protein kinase (MAPK) family are essential regulators of cell proliferation and differentiation (18–21). The *Toxoplasma* genome encodes three MAPKs: ERK7, MAPKL1, and MAPK2 (Figure 1A). ERK7 is conserved throughout eukaryota and we have recently shown that TgERK7 is essential for conoid biogenesis (22). TgMAPKL1 is found only in coccidian parasites and plays a role in preventing centrosome overduplication in order to ensure proper binary division through endodyogeny (3, 23). We have identified the cellular function of the third kinase, MAPK2, which is specific to and conserved throughout Alveolata. To uncover the function of MAPK2 in *Toxoplasma*, we applied the auxin-inducible degron (AID) system to inducibly degrade the protein. We found that parasites lacking TgMAPK2 arrested early in cell cycle, which eventually led to parasite death. While these parasites failed to duplicate their centrosomes, and never initiated daughter cell budding, they continued to grow in size and to replicate their Golgi, mitochondria, and apicoplasts. Our data implicate TgMAPK2 as an essential regulator of an early check point that is required to couple *Toxoplasma* cell growth with completion of the cell cycle.

**Figure 1.**
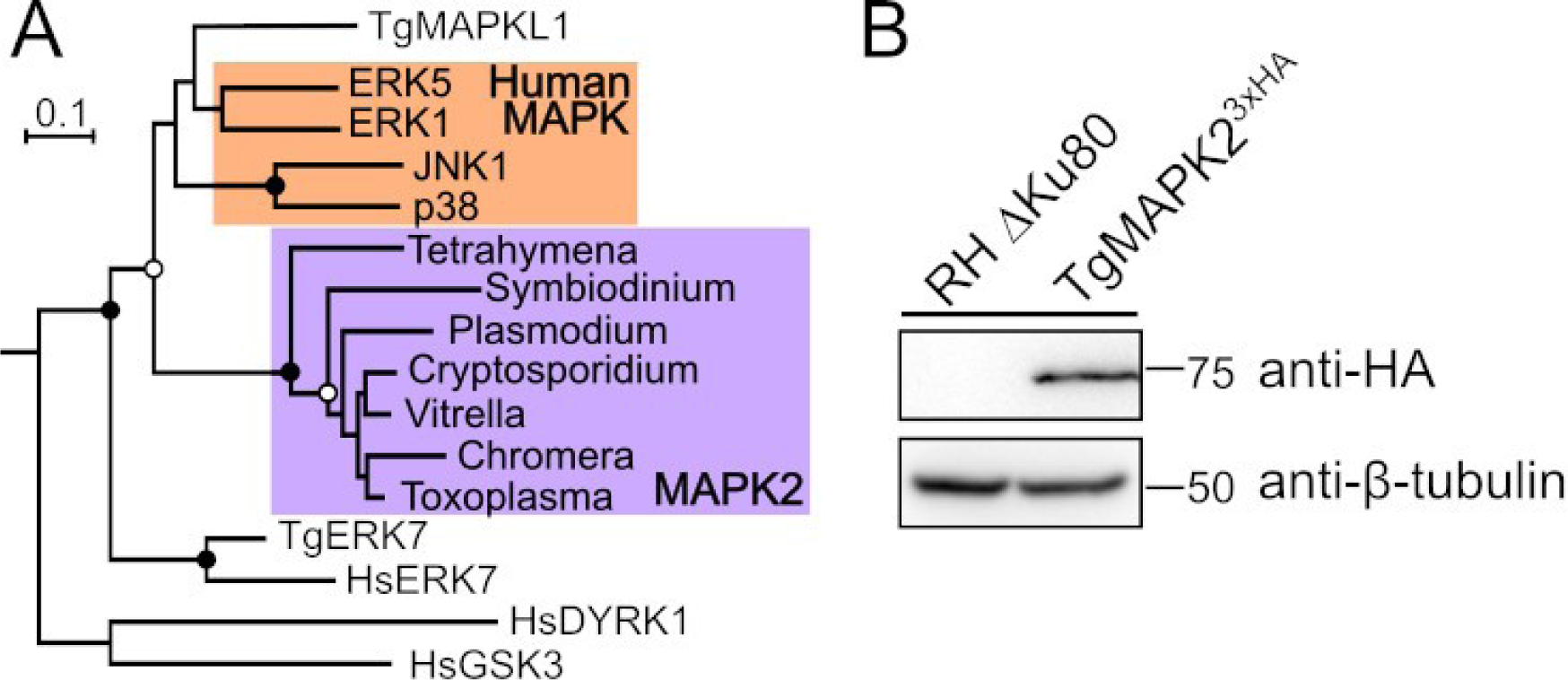
(A) Phylogenetic tree demonstrates MAPK2 represents a distinct MAPK subfamily (highlighted purple) that is conserved throughout Alveolata. Human MAPKs ERK1, ERK5, JNK1, and p38 are highlighted orange. Human DYRK1, GSK3, and ERK7 and *Toxoplasma* MAPKL1 and ERK7 were used as outgroups. Bootstrap support indicated by: Black circles >95%; Open circles >80%. (B) Western blot on TgMAPKL2^3xHA^ and parental lysates. β-Tubulin (anti-β-Tubulin) was used as a loading control.

## Results

### MAPK2 localizes to cytoplasmic puncta in Toxoplasma

The *Toxoplasma* genome encodes three MAPKs. Two of these kinases have been characterized: TgMAPKL1 (TGME49_312570) prevents centrosome overduplication (3) and TgERK7 (TGME49_233010) is required for conoid biogenesis (22, 24). The gene TGME49_207820 encodes the *Toxoplasma* ortholog of MAPK2, a MAPK that is specific to and conserved in all Alveolates, suggesting a specialized function (Figure 1A). We first sought to determine the subcellular localization of TgMAPK2. To this end, we engineered a parasite strain to express the native TgMAPK2 with a C-terminal 3xHA epitope tag (TgMAPK2^3xHA^) using a CRISPR-mediated double homologous recombination strategy. A western blot of TgMAPK2^3xHA^ parasite lysate stained with anti-HA antibody showed a single band of the expected mass (66-kD; Figure 1B).

Our initial immunofluorescence analysis (IFA) revealed that TgMAPK2 appears as puncta dispersed throughout the parasite cytosol (Figure 2A). We therefore co-stained with anti-Tgβ-tubulin and several other well-characterized markers for parasite organelles. We observed no co-localization of TgMAPK2^3xHA^ puncta with any organellar markers we tested (Figure 2A,B), including β-tubulin, Hoechst, mitochondrion (anti-TOM40), apicoplast (anti-ACP), rhoptries (anti-ROP2), or centrosome/conoid (Centrin 1 and Centrin 2). We reasoned that the TgMAPK2 puncta may represent intracellular vesicles and went on to test its co-localization with markers for intracellular trafficking including the Golgi (GRASP55) and endolysosomal trafficking system (*Toxoplasma* Rab proteins) (25). While many of these markers also appear punctate, confocal imaging revealed TgMAPK2^3xHA^ does not localize to structures stained by any of GRASP55, Rab5a, Rab6, or Rab7 (Figure 2A,B). TgMAPK2 therefore appears to mark a structure that is distinct from well-characterized organelles and trafficking machinery in *Toxoplasma*. Moreover, we observed that the TgMAPK2 signal dropped to near undetectable levels late in the parasite cell cycle (Figure 2C). While this loss of signal intensity may be due to dissolution of the TgMAPK2 puncta, it is consistent the variation of TgMAPK2 transcript levels through the parasite cell cycle, with a minimum during mitosis (Figure 2D; (26)).

**Figure 2.**
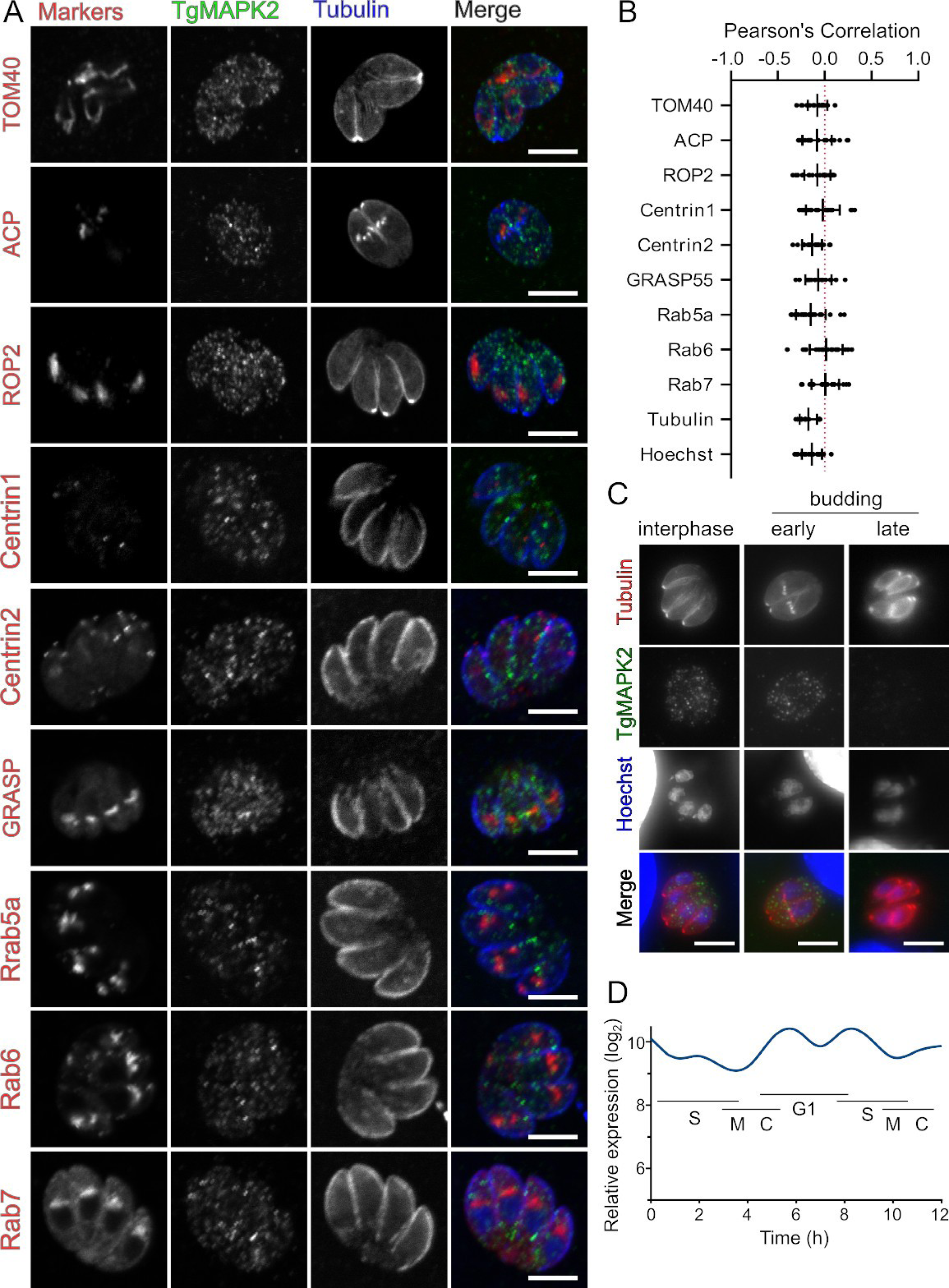
TgMAPK2 forms cytosolic puncta in interphase and early budding parasites. (A) 0.5-µm confocal slices of TgMAPK2^3xHA^ and GFP-α-Tubulin (blue) expressing parasites were either stained or transiently transfected (Centrin and Rab FP fusions) with the indicated markers. (B) Pearson’s correlation coefficients for TgMAPK2^3xHA^ and the indicated markers from (A); n=∼20 cells. (C) TgMAPK2^3xHA^ intensity diminishes late in the cell cycle. (D) Transcript levels of MAPK2 peak during G1/early S (data (26), retrieved from ToxoDBv43). All scale bars: 5 µm.

We next used subcellular fractionation to attempt to determine whether TgMAPK2 behaved as a soluble or has appreciable membrane or cytoskeletal interaction. TgMAPK2^3xHA^ parasites were lysed by freeze-thaw and ultracentrifiged to separate soluble (HSS) and insoluble (HSP) fractions. The HSP was resuspended in either PBS, 0.1 M Na_2_CO_3_, pH 11.5 (high pH), 1 M NaCl (high salt) or 1% TritonX-100 (TX100) and incubated at 4°C for 30 minutes, followed again by ultracentrifugation. We found that while the vast majority of the TgMAPK2 protein in the first HSS fraction, a small portion of the protein was in the HSP, and was only soluble in detergent (Supplemental Figure S1). Thus TgMAPK2 behaves as a soluble protein, though a small portion of the protein (<5%) may be integrally associated with a cellular membrane.

### TgMAPK2 is essential for the completion of parasite lytic cycle

We were unable to obtain MAPK2 knockouts using either homologous recombination or CRISPR-mediated strategies, so we applied the AID system. The AID system allows the conditional degradation of target proteins upon addition of a small molecule auxin (indole-3-acetic acid; IAA), and has been recently adapted to *Toxoplasma* (27). We engineered a parasite strain in which the MAPK2 protein was expressed in-frame with an AID and 3xFLAG tag at the C-terminus in the background of RHΔ*ku80* expressing the rice TIR1 auxin response protein (TgMAPK2^AID^). TgMAPK2 localization appeared unaffected by the addition of the AID tag, and the TgMAPK2^AID^ was degraded upon addition of 500 µM auxin indole-3-acetic acid (IAA) (Figure 3A). TgMAPK2 protein was undetectable by western-blot after 15 minutes of IAA treatment (Figure 3B). We will refer to parasites in which TgMAPK2 has been inducibly degraded as TgMAPK2^AID/IAA^. While TgMAPK2^AID^ parasites produced normal plaque numbers, TgMAPK2^AID/IAA^ parasites produced no plaques (Figure 3C). To confirm that this phenotype is due to the absence of TgMAPK2 protein, and to determine whether TgMAPK2 kinase activity is required for its function, we expressed an extra, non-degradable copy of wild-type (WT) or kinase-dead (KD; D271A) kinase in the background of TgMAPK2^AID^ parasites (Figure 3D). While the WT-complemented TgMAPK2^AID^ parasites were able to form plaques even in the presence of IAA, the KD-complemented parasites were not (Figure 3E). TgMAPK2 kinase activity therefore appears required for its function. We note that WT-complemented parasites did not entirely rescue plaque formation. This is likely due to two factors: (1) we were unable to obtain stable clones of ectopically expressed MAPK2 driven by its native promoter, and instead used the dihydroflate reducatase (DHFR) promoter; (2) ectopic MAPK2 expression appeared mosaic in a clonal population; 50-70% of parasites expressed the complemented copy at any given time, consistent with our rescue data. Taken together, the above data clearly demonstrate that TgMAPK2 is essential for the lytic cycle, which is consistent with our inability to genetically disrupt the TgMAPK2 gene.

**Figure 3.**
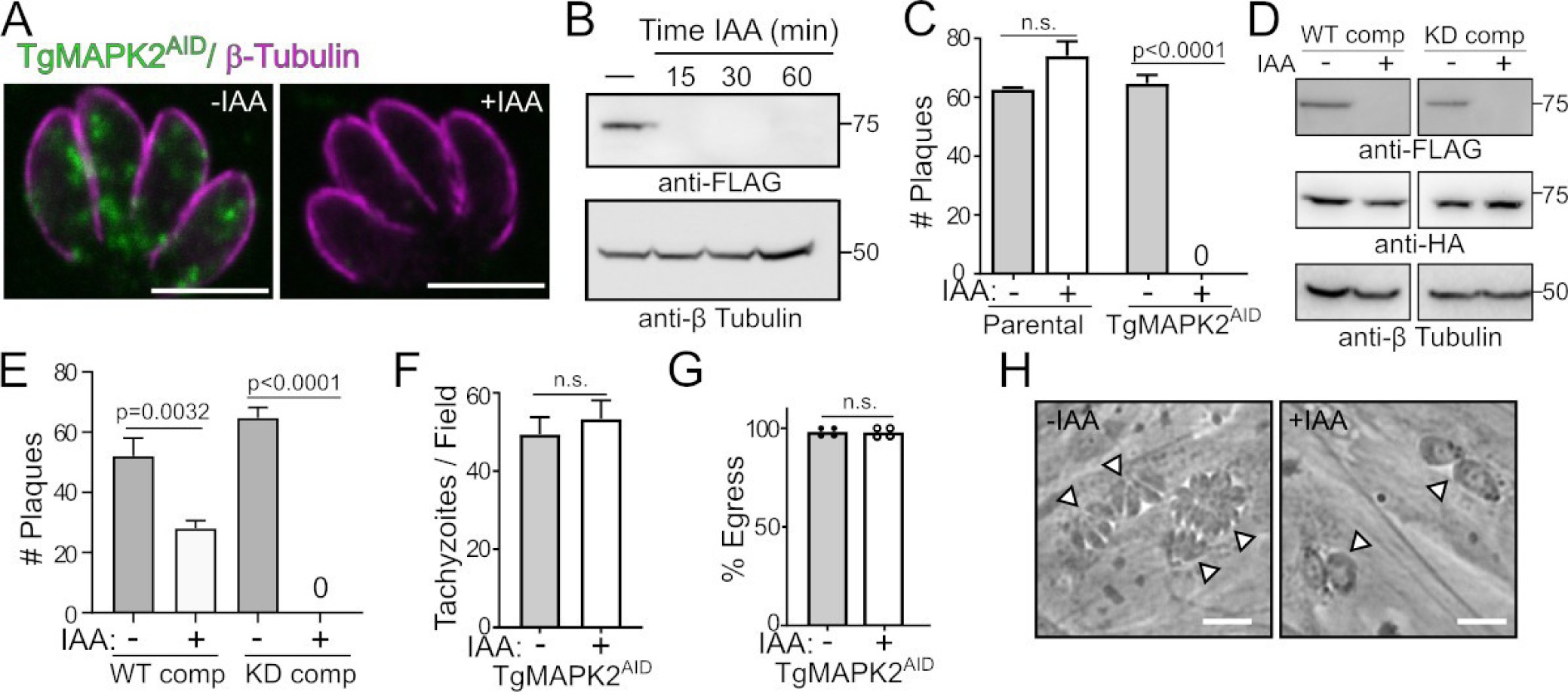
TgMAPK2 is essential for parasite proliferation. (A) 0.5-µm confocal slice of TgMAPK2^AID^ parasites using anti-FLAG (green) and anti-Tgβ-tubulin (magenta) in the presence and absence of IAA. Scale bar = 5 µm. (B) Western blot of TgMAPK2^AID^ protein levels at increasing growth time in 500 μM IAA. Anti-Tgβ-tubulin was used as a loading control. (C) Quantification of triplicate plaque assays comparing parental and TgMAPK2^AID^ parasites grown in the presence and absence of IAA. (D) Protein levels of WT/KD complemtented TgMAPK2^3xHA^ are not effected by IAA and do not affect degradation of TgMAPK2^AID^. Anti-Tgβ-tubulin was used as a loading control. (E) Quantification of triplicate plaque assays comparing wild-type or kinase-dead complement parasites grown in the presence and absence of IAA. (F) invasion and (G) egress of TgMAPK2 parasites grown without and for the last 2 h with IAA. (H) Phase contrast micrograph comparing 20 h growth of TgMAPK^AID^ (-/+) IAA treatment. Arrowheads indicate individual vacuoles. Scale bars = 10 µm. p values are from two-tailed unpaired Student’s t-test.

Plaque assays report on the entire lytic cycle, comprising multiple rounds of invasion, replication, and egress. Degradation of TgMAPK2 did not significantly affect parasite invasion or egress from host cells (Figure 3F, 3G). TgMAPK2^AID/IAA^ parasites did not, however, replicate normally. Instead, we observed that many TgMAPK2^AID/IAA^ parasites were only able to undergo a single round of replication following treatment with 500 µM IAA for 20 hours, after which they arrested and showed an aberrant morphology under phase contrast microscopy (Figure 3H). While TgMAPK2^AID^ parasites replicated normally, more than 80% of the TgMAPK2^AID/IAA^ vacuoles contained two enlarged, morphologically aberrant parasites. The remaining TgMAPK2^AID/IAA^ vacuoles contained a single oversized parasite (Figure 3H). These data led us to hypothesize that TgMAPK2 plays a crucial role at an early stage in parasite division.

### TgMAPK2 knock-down arrests parasites prior to initiation of daughter cell budding

During acute infection, the *Toxoplasma* tachyzoite replicates by endodyogeny, in which two daughter cells are assembled within the mother (6). This mechanism of division requires that organellar biogenesis be tightly coupled to the cell cycle. The parasite cellular ultrastructure thus changes drastically as cell cycle progresses, which provides us with a number of markers that can easily distinguish cell cycle stages via fluorescent microscopy and electron microscopy.

In a normal culture, the *Toxoplasma* cell cycle is synchronized within individual vacuoles (28), but asynchronous among vacuoles in a population, even among parasites that have invaded at the same time. Because parasites divide asynchronously, at any given time the parasites in a population occupy all stages of the cell cycle. We therefore reasoned we could identify the point of arrest upon TgMAPK2 degradation by examining arrested parasites and determining which structures are lost during increasing times of growth in the presence of IAA. Here we used a set of fluorescent markers to classify the parasite cell cycle into three broad categories: (i) “Before budding”, (ii) “Budding” and (iii) “Cytokinesis” (Figure 4A). To distinguish these categories, we chose as markers (i) the *Toxoplasma* inner membrane complex 1 (IMC1) which stains the mother outline and growing daughter scaffolds; (ii) β-Tubulin, which labels the subpellicular microtubules and conoids; and (iii) Hoechst, which labels nuclear and plastid DNA. We engineered the TgMAPK2^AID^ strain to co-express IMC1-mVenus and mTFP1-α-tubulin. TgMAPK2^AID^ parasites were allowed to invade a fresh monolayer of fibroblasts for 2 hours, after which uninvaded parasites were washed off. Media were changed to +IAA after increasing 2 hour increments, and all parasites were grown for a total of 10 hours (Figure 4A,B). Parasites were then fixed and prepared for imaging.

**Figure 4.**
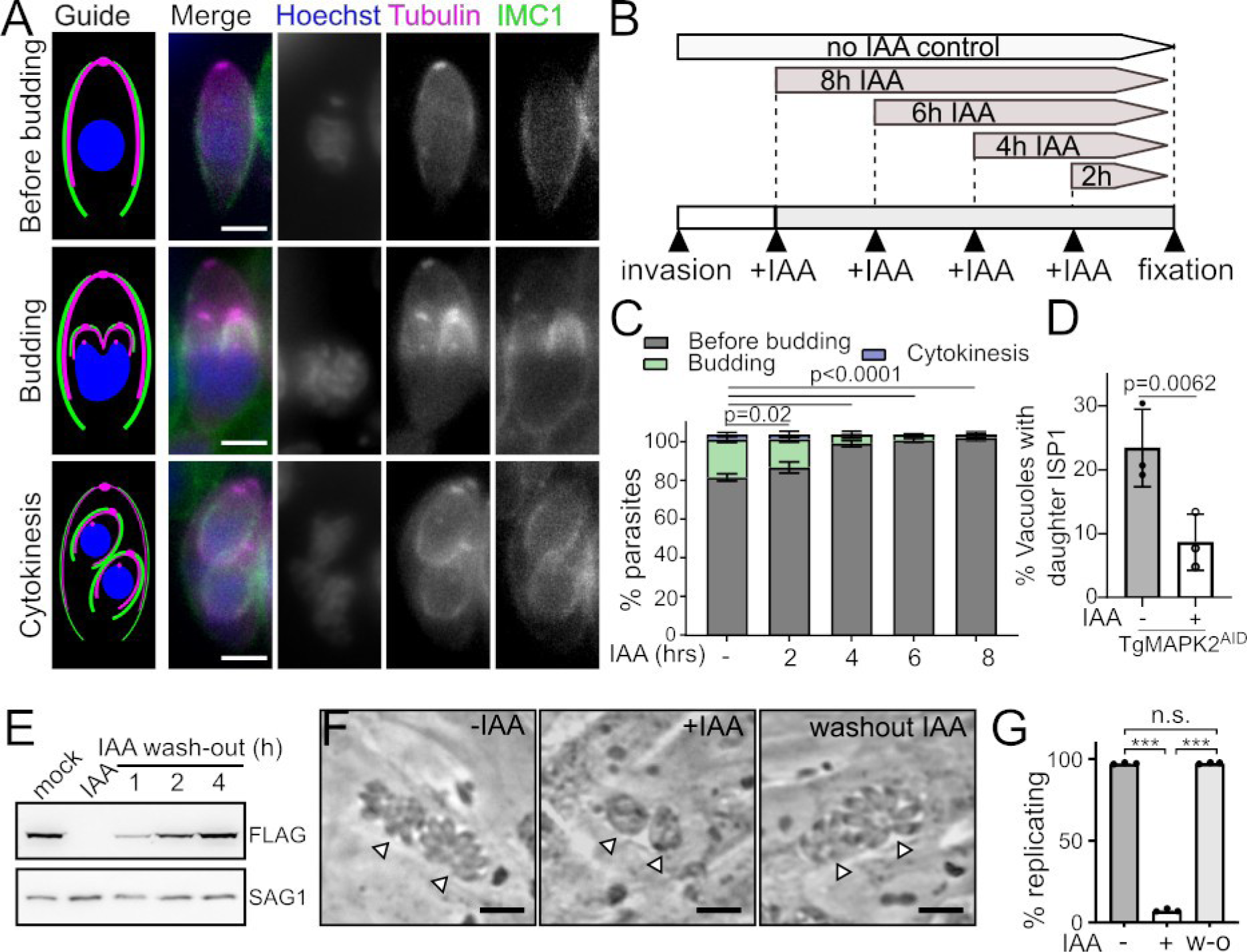
Loss of TgMAPK2 leads to a defect in daughter cell budding. TgMAPK2^AID^ parasites stably expressing mTFP1-α-Tubulin (magenta) and TgIMC1-mVenus (green) were stained with Hoechst 33342 (blue) to distinguish the three indicated categories in cell cycle. Scale bars: 2 µm. (B) Experimental flow of addition of IAA in increasing 2h increments. (C) Quantification of parasites in each category with increasing time of growth in IAA. Mean±SD of n=3 biological replicates; 200-300 parasites counted per condition. (D) Quantification of parasites with daughter cell budding ring (ISP1 in daughters buds) grown in the presence or absence of IAA for 8 hours. Mean±SD of n=3 biological replicates; 200-300 parasites counted per condition. p values are from two-tailed Student’s t-test. (E) Western blot demonstrates MAPK2^AID^ protein levels are restored 2-4 h after IAA washout. (F) Phase contrast micrograph comparing growth of TgMAPK2^AID^ parasites over 18 h in IAA or after 2 h in IAA with an additional 16 h after wash-out. Arrowheads indicate individual vacuoles. Scale bars = 10 µm. Quantification of percent of vacuoles that appeared to be replicating normally (≥4 parasites/vacuole) of n=3 independent replicates as in (G). p values are from one-way ANOVA with Dunnett’s test. (p<0.0001: ***).

We quantified the number of parasites in each rough stage of the cell cycle for increasing incubation time in +IAA media (Figure 4C). We observed that parasites grown without IAA had 79±2% “before budding,” 19±2% budding, and 3±1% parasites undergoing cytokinesis. As time in IAA increased, we observed a marked decrease in the number of parasites that were budding and undergoing cytokinesis. After of 6-8 hours of IAA treatment, 98±2% of the parasites were in a non-budding state. Taken together, these data demonstrate that the arrest we observe due to degradation of TgMAPK2 is prior to initiation of daughter cell budding, after which TgMAPK2 does not appear essential for division. To verify the defect in the initiation of daughter cell budding in the absence of TgMAPK2, we tested IMC Sub-compartment Protein 1 (ISP1) labeling of parasites, which is an early marker for daughter bud formation (29). Consistent with the quantification of IMC1 staining (Figure 4C), ISP1-staining demonstrated daughter cell budding rings appeared in 23±6% of the parasites without IAA, whereas only 8±4% parasites had daughter budding rings after 8 hours of IAA treatment (Figure 4D).

A regulatory arrest in cell cycle should be reversible by restoration of the protein. Induced degradation of an AID-tagged protein can be reversed by washing-out IAA (Figure 4E and (30)). We therefore asked whether the block we had observed in parasite division due to TgMAPK2 degradation was reversible. TgMAPK2^AID^ parasites were allowed to invade host fibroblasts, and then grown for 8 h in the presence or absence of IAA. Parasites were then allowed to either continue to grow in IAA for an additional 18 h, or the media changed to -IAA before continued growth. Parasites were then fixed and imaged by phase contrast microscopy. As expected, TgMAPK2^AID^ parasites grown continuously in IAA were arrested. Parasites that had been grown for only 8 h in IAA before wash-out, however, were indistinguishable from those that had been grown without IAA (Figure 4F,G). Thus, the block in *Toxoplasma* division caused by TgMAPK2 degradation likely represents a checkpoint arrest.

### TgMAPK2 *degradation impairs centrosome duplication*

An early, crucial event prior to *Toxoplasma* daughter cell budding is the duplication of the centrosome (8). Centriole duplication during the G1/S boundary of the cell cycle marks entry into the S phase and provides a spatial direction for the assembly of the daughter buds (31). Earlier work identified an unusual bipartite structure of the centrosome in *Toxoplasma* which includes two cores: the outer core, distal from nucleus, and the inner core, proximal to nucleus (3). These cores have distinct protein compositions, by which they can be easily distinguished (3). We therefore checked whether the duplication of either centrosomal core was compromised in TgMAPK2^AID/IAA^ parasites. We engineered the TgMAPK2^AID^ strain to stably express an mVenus fusion to the centrosomal outer core protein TgCentrin1, and a 3xHA fusion to the centrosomal inner core protein TgCEP250L1. Freshly invaded parasites were grown in the presence or absence of IAA for 8 hours. Anti-HA was used to stain TgCEP250L1, and Hoechst 33342 was included to track the nuclear DNA (Figure 5A). We quantified the percentage of parasites with two clearly separated centrosomes based on both inner core marker TgCEP250L1 and outer core marker mVenus-TgCentrin1 signals. As expected, ∼40% of TgMAPK2^AID^ parasites grown in the absence of IAA had successfully duplicated both their centrosomal inner cores and outer cores (Figure 5B,C), consistent with published reports (5, 8). In contrast, depletion of MAPK2 profoundly suppressed duplication of both cores, though the phenotype was less severe for the inner core than the outer core (Figure 5B,C).

**Figure 5.**
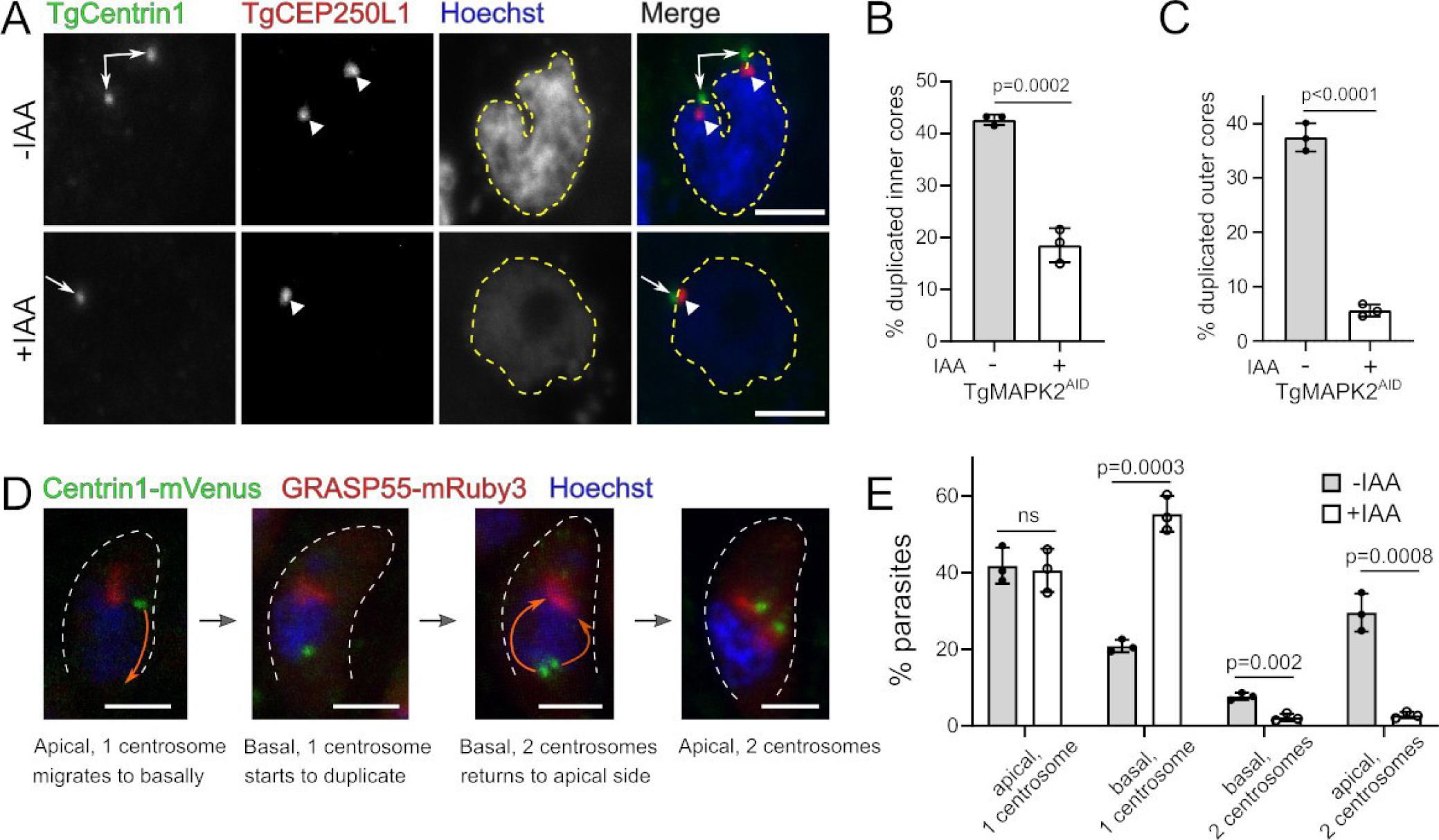
TgMAPK2 degradation impairs centrosome duplication after basal migration. (A) Z-projections of confocal stacks of TgMAPK2^AID^ parasites stably expressing mVenus-TgCentrin1 (green) and TgCEP250L1-3xHA (red) were grown for 8 hours in the presence (+) or absence (-) of 500 µM IAA, and co-stained with Hoechst (blue). TgCentrin1 (arrows), TgCEP250L1 (large arrowheads). Nuclei are outlines in yellow. (B) Quantification of parasites with duplicated centrosomal inner cores (TgCEP250L1) (B) and outer cores (TgCentrin1) (C) grown in the presence or absence of IAA for 8 hours. Mean±SD of n=3 biological replicates; 200-500 parasites counted per condition. (D) maximum intensity Z-projections of confocal stacks of TgMAPK2^AID^ parasites stably expressing mVenus-TgCentrin1 (green) and GRASP55-mRuby3 (red) during the cell cycle; representative images summarizing centrosome migration/duplication are shown.. (E) Quantification of parasites in each category shown in (D) grown in the presence or absence of IAA for 8 hours. Mean±SD of n=3 biological replicates; 300-500 parasites counted per condition. p values are from two-tailed Student’s t-test. All scale bars: 2 µm.

During late G1 phase, the centrosome migrates to the basal end of the nucleus, where it duplicates and then separates. Subsequently, duplicated centrosomes traffic back to the apical end of the nucleus, where they re-associate with the Golgi (32). To check whether the non-duplicated centrosome migrates to the basal end, we engineered the TgMAPK2^AID^ strain to express an mVenus fusion to the centrosome marker TgCentrin1, and an mRuby3 fusion to the Golgi marker GRASP55. Freshly invaded parasites were grown in the presence or absence of IAA for 8 hours, and Hoechst 33342 was included to track the nuclear DNA. We summarize the centrosome migration/duplication cycle into four steps: (i) “apical, 1 centrosome”, (ii) “basal, 1 centrosome”, (iii) “basal, 2 centrosomes” and (iv) “apical, 2 centrosomes” as shown in Figure 5D. We quantified the number of parasites in each step in the presence or absence of IAA. As expected, we found that treatment with IAA drastically reduced the number of duplicated centrosomes (Figure 5E). Intriguingly, IAA treatment resulted in a ∼2.6-fold increase (55.3±5% vs. 21±2%) of parasites in which the centrosome had migrated to the basal end of the parasite but failed to duplicate. Therefore, we conclude that TgMAPK2 depletion does not affect centrosome migration to the basal end of the nucleus, while it does compromise duplication of the centrosome.

### Knock-down of TgMAPK2 results in a defect in DNA replication impairs entry into mitosis

DNA replication and nuclear division are crucial for each daughter cell to inherit one copy of the cell’s genetic material. We observed substantially weaker nuclear Hoechst staining in TgMAPK2^AID/IAA^ than in control parasites when imaged at the same microscope settings (Figure 5A). As we observed this phenotype even after relatively short, 8h growth in IAA, the low-Hoechst intensity phenotype occurs early after loss of TgMAPK2 and is not a secondary effect or artifact of prolonged growth in IAA. This phenotype suggests that DNA replication and karyokinesis may also be inhibited in TgMAPK2^AID/IAA^ parasites. To test this hypothesis, we stained the parasite DNA with DAPI and performed a FACS-based analysis to compare the DNA content in TgMAPK2^AID^ to parasites grown in 8h of IAA. 71±6% of control parasites possessed 1N DNA content, while 29±6% of parasites possessed 1.8∼2N DNA content (Figure 6A). In TgMAPK2^AID/IAA^ parasites, however, the majority (55±3%) of parasites possess >1N DNA content, though the peak is left-shifted (Figure 6B), indicating that TgMAPK2^AID/IAA^ parasites can initiate, but not complete, DNA replication. Thus, TgMAPK2^AID/IAA^ parasites are unable to progress successfully through S phase.

**Figure 6.**
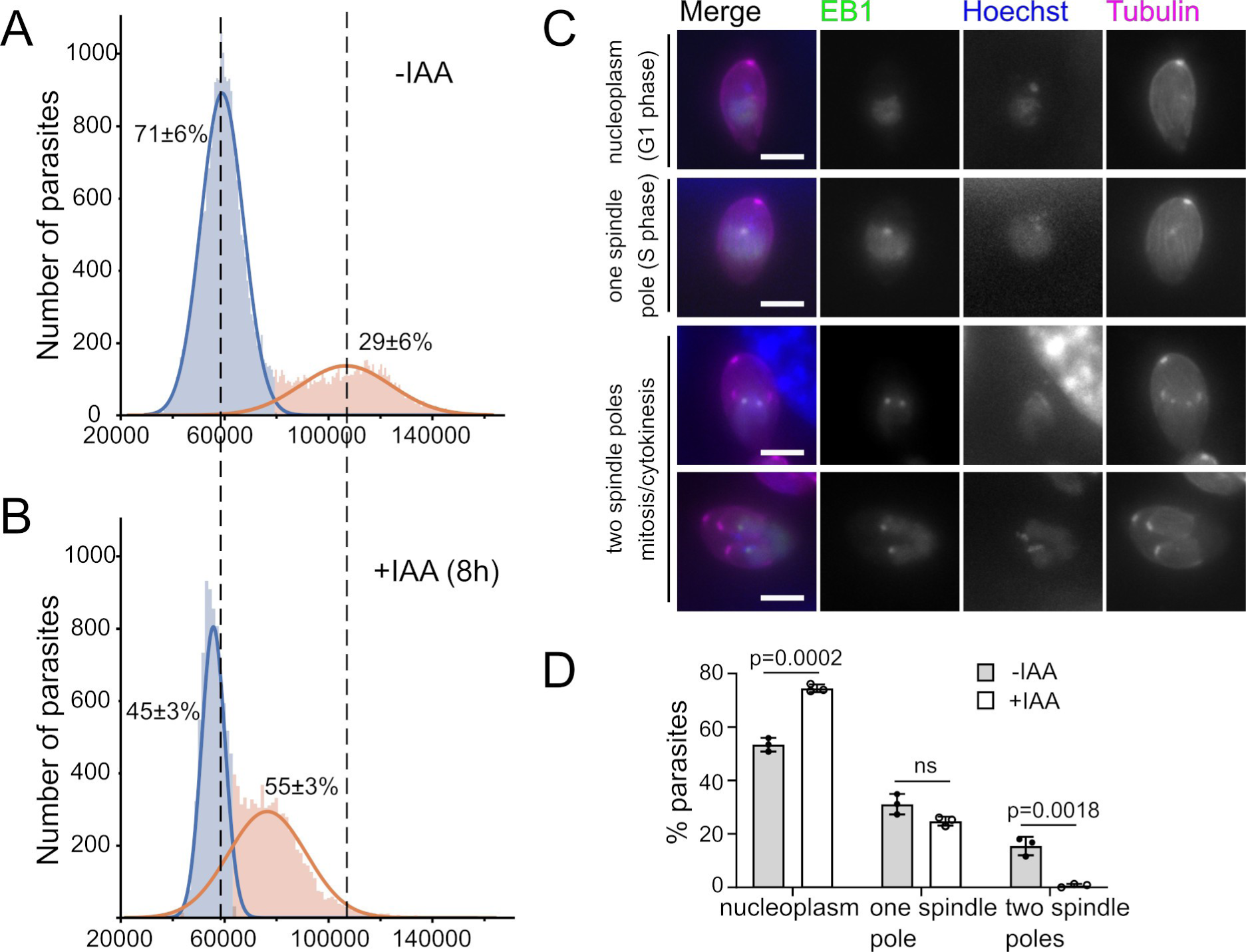
TgMAPK2 degradation results in DNA replication defect and impairs mitosis. (A-B) Histogram showing DNA content were measured by FACS analysis. TgMAPK2^AID^ parasites growing in the presence or absence of IAA for the last 8 hours were labeled with 1 µg/ml DAPI, and 50,000 events were collected from each of the three replicates. Gaussian mixtures were estimated for individual FACS run, and area under gaussian curves was measured to quantify. (C) TgMAPK2^AID^ parasites stably expressing TgEB1-mRuby3 (green) and mTFP1-α-tubulin (magenta) were imaged, together with Hoechst staining. Representative images of the each stage are shown. Scale bars: 2 µm. (D) Quantification of parasites in each stage shown in (C) grown in the presence or absence of IAA for 8 hours. Mean±SD of n=3 biological replicates; 80-200 parasites counted per condition. p values are from unpaired two-tailed Student’s t-test.

The coordination of the spindle assembly and the centrosome cycle are critical for mitosis progression. During the *Toxoplasma* cell cycle, formation of spindle poles is coincident with completion of DNA replication and nucleus lobulation (14, 33). In order to determine whether mitosis initiates in TgMAPK2^AID/IAA^ parasites, we used the microtubule end-binding protein 1 (EB1) as a marker. EB1 distributes in the nucleoplasm during interphase, but transitions to the spindle poles when mitosis starts (33). The TgMAPK2^AID^ strain was engineered to stably express TgEB1-mRuby3. Freshly invaded parasites were grown in the presence or absence of IAA for 8 hours, and Hoechst 33342 and β-tubulin were then included to track the nuclear DNA and mark the parasite boundary. We classified the localization of TgEB1 into (i) “nucleoplasm” (interphase), (ii) “one spindle pole” (S phase), and (iii) “two spindle poles” (mitosis/cytokinesis) (Figure 6C). Next, we counted the parasite numbers in each class based on the TgEB1-mRuby3 signal. We observed that without IAA treatment, 53±3% of TgMAPK2^AID^ parasites presented nucleoplasm TgEB1 localization, as compared to 75±1% in TgMAPK2^AID/IAA^ parasites. 31±4 of TgMAPK2^AID^ parasites and 25±2% of TgMAPK2^AID/IAA^ parasites displayed one spindle pole, indicating successful entry into S-phase. However, while 16±3% of TgMAPK2^AID^ parasites showed two spindle poles of TgEB1 localization, essentially none of the TgMAPK2^AID/IAA^ parasites did (Figure 6D). Taken together, our results indicate that TgMAPK2^AID/IAA^ parasites are able to enter S-phase, where they arrest, unable to proceed into mitosis.

### Degradation of TgMAPK2 does not block replication of mitochondrion, apicoplast, or Golgi apparatus

To better understand the nature of the arrest upon TgMAPK2 depletion, we next sought to characterize the morphological changes to the parasite after this arrest. We used transmission electron microscopy (TEM) to compare the ultrastructure of TgMAPK2^AID^ parasites grown in the absence or presence of IAA for a short time (6 hours), which was sufficient to induce arrest. We observed well-formed daughter buds in ∼20% of the asynchronously dividing TgMAPK2^AID^ parasites (Figure 5A), which is consistent with our IFA quantification (Figure 4). While the TgMAPK2^AID/IAA^ parasites did not show obvious ultrastructural defects, daughter cell budding appeared completely blocked (Figure 7C), again consistent with our IFA quantification. We next examined parasites that were incubated for extended periods in IAA. After 20 hours in IAA, the architecture of TgMAPK2^AID/IAA^ parasites was misshapen, as evidenced by enlarged cells and loss of the parasite’s distinctive shape (Figures 7E-H). In addition, we observed an over-duplication of many organelles, including the Golgi apparatus and apicoplasts (Figures 7E-H). Moreover, at these longer time points, we observed vacuoles containing enlarged residual bodies filled with organellar debris, even in vacuoles containing only a single (albeit misshapen) parasite (Figures 7E).

**Figure 7.**
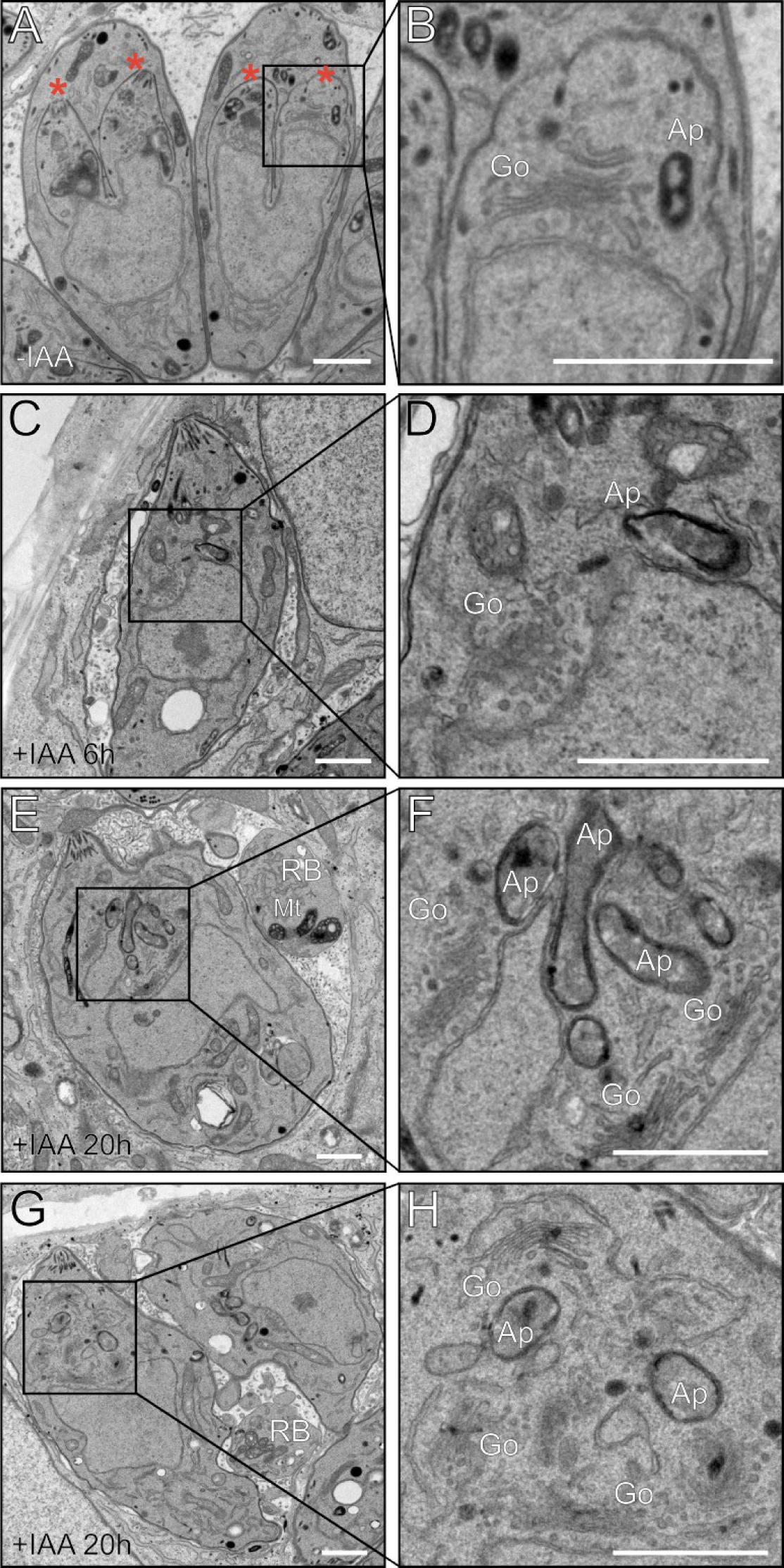
Prolonged incubation in IAA results in multiple defects in TgMAPK2^AID^ parasite ultrastructure. Transmission electron micrographs of intracellular TgMAPK2^AID^ parasites in (A,B) the absence of IAA, 6 h growth in IAA (C,D) or 20 h IAA (E,F and G,H). Daughter buds (*), Golgi apparatus (Go), Apicoplast (Ap), Mitochondria (Mt); Residual bodies (RB) are marked. All scale bars: 1 µm. Note that 100% of parasites treated for 20 h IAA had overduplicated apicoplasts, and 35±5% had overduplicated Golgi; neither phenotype was observed in untreated parasites.

In order to scrutinize the organelle replication progress and centrosome duplication simultaneously in TgMAPK2^AID/IAA^ parasites, mVenus-Centrin1 expressing parasites were stained with antibodies recognizing either TOM40 (mitochondrion) or ACP1 (apicoplast). Individual parasites and their daughter buds were identified by IMC1 staining. Parasites were grown for 20h in the presence or absence of IAA. Our microscopic analysis confirmed that TgMAPK2^AID^ parasites replicated normally and maintained a normal count of apicoplast and mitochondrion per parasite (Figure 8A,B). Strikingly, in TgMAPK2^AID/IAA^ parasites, mitochondria and apicoplasts continued to grow even after arrest, though without centrosome duplication and new daughter cell buds, these organelles failed to separate and partition (Figure 8A,B).

**Figure 8.**
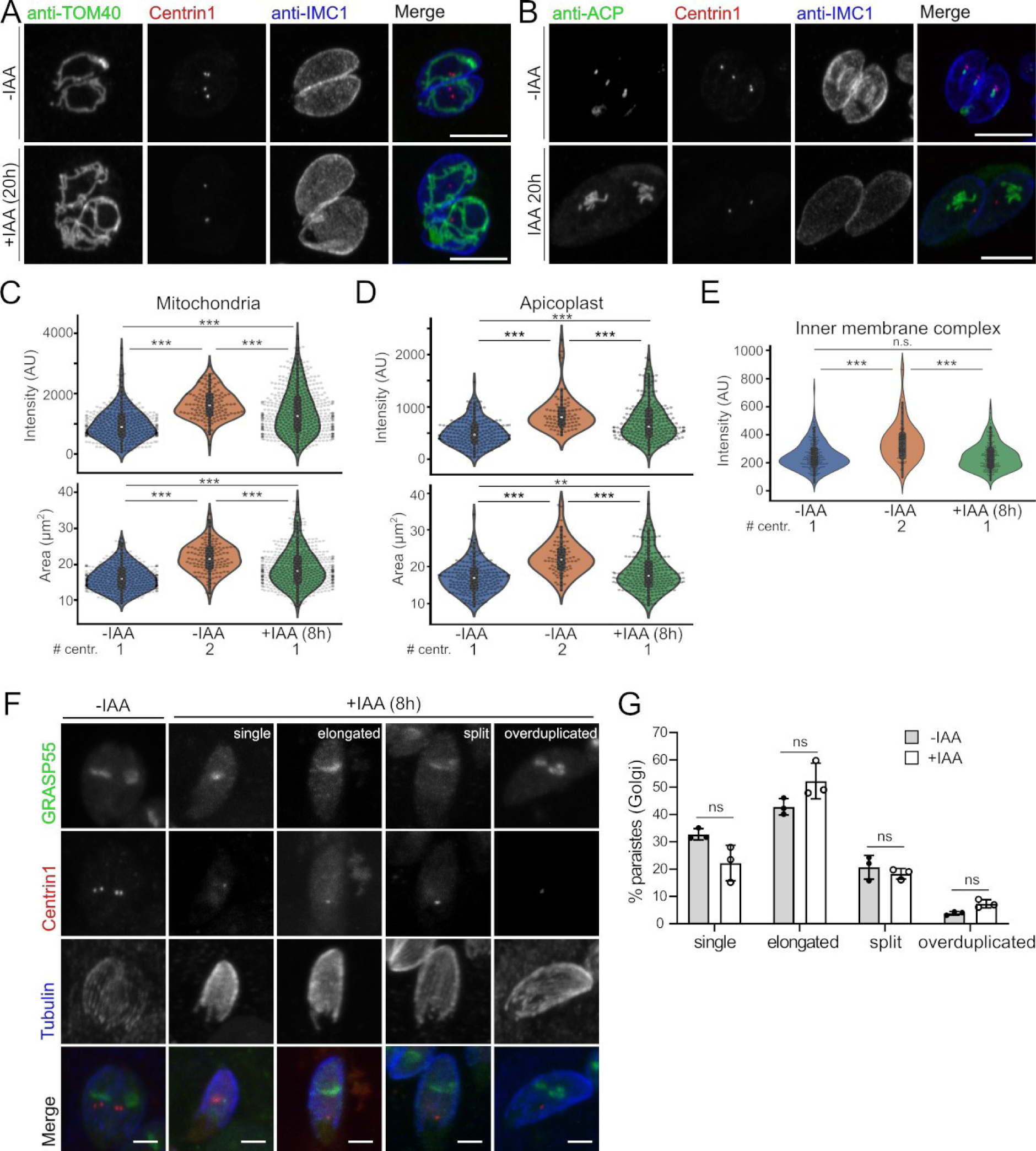
Loss of TgMAPK2 does not block organelle replication. (A-B) Z-projections of confocal stacks of the TgMAPK2^AID^ parasites stably expressing mVenus-TgCentrin1 (red) grown for 20 hours in the presence or absence of IAA. Parasites were fixed and stained with antibodies for IMC1 (blue), TOM40 (green), or ACP1 (green). Scale bars: 5 µm. (C-E) The total intensities of TOM40 (mitochondria), ACP (apicoplasts), and IMC1 as well as the areas of the same set of parasites were measured and quantified in each of the class. (F) Z-projections of confocal stacks of the TgMAPK2^AID^ parasites stably expressing mVenus-TgCentrin1 (red) and TgGRASP55-mRuby3 grown in the presence or absence of IAA for 8 hours, and co-stained with anti-Tgβ-tubulin. The representative images of each class presenting Golgi morphologies were shown. (G) Quantification of parasites in each class shown in (F) grown in the presence or absence of IAA for 8 hours. Mean±SD of n=3 biological replicates; 200-300 parasites counted per condition. p values are from two-tailed Student’s t-test.

We next asked whether the increased apicoplast and mitochondrial load we observed after 20h growth in IAA was also observable at earlier time points. Using the same staining strategy as above, parasites were grown in the presence or absence of IAA for 8h before analysis. As >95% of TgMAPK2^AID/IAA^ parasites possessed a single centrosome at this time point in our dataset, we focused on this majority population. We quantified the intensity of organellar staining as a proxy for organelle load in both control and IAA-treated parasites. As expected, control parasites with duplicated centrosomes had increased mitochondria and apicoplast load as compared to cells with a single centrosome. TgMAPK2^AID/IAA^ parasites, however, displayed a non-normal distribution of mitochondrial and apicoplast load that was significantly different from both control populations (Fig 8C,D). Using the IMC1 signal as a guide, we quantified the size of each parasite. As expected, control parasites that had initiated budding were consistently larger than single-centrosome parasites (Fig 8C,D). TgMAPK2^AID/IAA^ parasites, however, had a complex distribution of sizes that was again distinct from both control populations.

All organisms’ cells match their organellar load to their need, which is quite often correlated with the cell’s size. We found that for all populations observed above, cell size was correlated with apicoplast and mitochondrial load (Supplemental Figure S2). As a control, we reanalyzed the ACP1-stained dataset, quantifying IMC1-levels (Figure 8E). Consistent with the idea that the IMC is created during cell division (34), rather than throughout growth, we found that control parasites with 2 centrosomes had higher levels of IMC1 than those with a single centrosome (Figure 8E). Importantly, IMC1 levels in IAA-treated cells were indistinguishable from control parasites with a single centrosome (Figure 8E), and IMC1 levels were constant across cell size in IAA-treated cells (Supplemental Figure S2).

Centrosome duplication and Golgi division are two of the earliest events in the highly organized parasite cell cycle (14). The centrosomes are always associated with Golgi except ∼1 hour of migration to the basal end of nucleus (14, 32). To examine the relationship between centrosome duplication and Golgi behaviors, we used TgMAPK2^AID^ parasites that stably expressing both mVenus-Centrin1 and the Golgi marker GRASP55-mRuby3. Parasites were grown with or without IAA for 8 hours, and β-tubulin was visualized to mark the parasite boundary. We binned and quantified the Golgi morphologies as (i) “single”, (ii) “elongated”, (iii) “split”, and (iv) “overduplicated” based on GRASP55-mRuby3 signal (Figure 8F). Our analysis showed that the Golgi appeared to replicate normally in TgMAPK2^AID/IAA^ parasites, even without centrosome duplication (Figure 8G).

Taken together, these data demonstrate that without TgMAPK2, the cell and organellar growth continues unhindered, and is therefore uncoupled from the parasite cell cycle.

### The two MAPKs, MAPKL1 and MAPK2, are required at different check points during the cell cycle

Our data show that TgMAPK2 is required for replication of both centrosomal cores. Intriguingly, loss-of-function of another member of the MAPK family, TgMAPKL1 also causes a defect in parasite centrosome replication (3). Notably, the previous study was performed using a temperature-sensitive mutant that necessitated studies at longer timescales (20 h at non-permissive temperature) and at which we observed secondary effects using the AID system with MAPK2. We therefore sought to compare the phenotypes caused by depletion of these two MAPKs both at 8h and 20h. To this end, we generated a TgMAPKL1^AID^ strain using the same strategy as we used for TgMAPK2. Freshly invaded TgMAPK2^AID^ or TgMAPKL1^AID^ parasites expressing mVenus-Centrin1 were grown in the presence of IAA for 8 or 20 hours. Parasites were fixed and stained with anti-Tgβ-tubulin. We found that after 20 hours with IAA, ∼30% of TgMAPK2^AID/IAA^ parasites possess >1 centrosome compared with ∼5% at 8 hours IAA treatment (Figure 9C,D). Consistent with the published data (3), and in contrast to the TgMAPK2^AID/IAA^ phenotype, we observed that 32.2±5.4% of TgMAPKL1^AID/IAA^ parasites over-duplicated their centrosomes (*i*.*e*., >2) after 8 hours of IAA treatment, and 76.7±4.8% of TgMAPKL1^AID/IAA^ parasites underwent centrosome overduplication after 20 hours of IAA treatment (Figure 9A,C). Thus, these two divergent MAPKs both regulate centrosome replication though they have opposing phenotypes, and are thus likely control different checkpoints in the *Toxoplasma* cell cycle.

**Figure 9.**
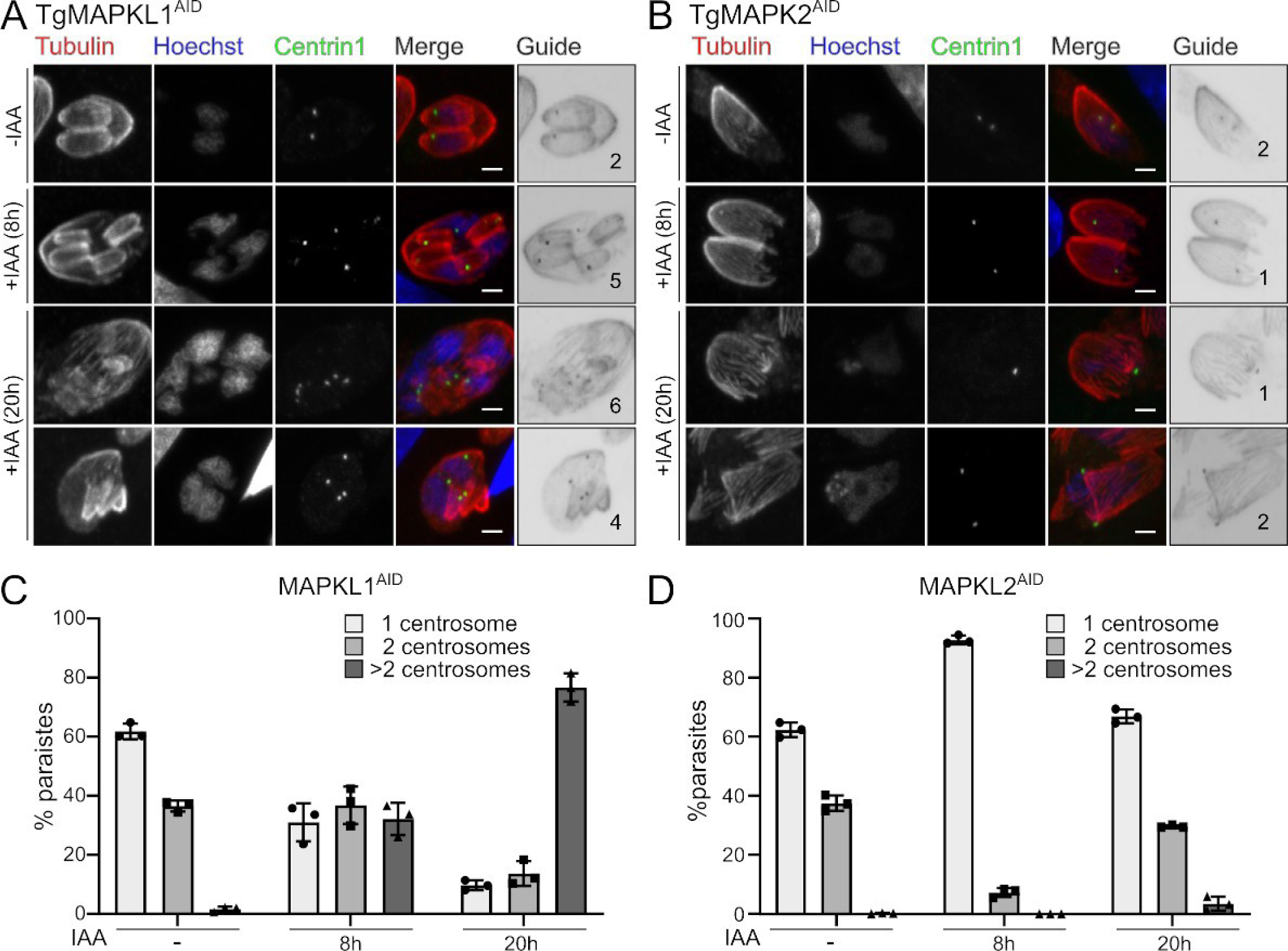
Centrosome duplication defect of TgMAPK2 is distinct from the phenotype caused by depletion of TgMAPKL1. (A-B) TgMAPKL1^AID^ or TgMAPK2^AID^ parasites expressing TgCentrin1-mVenus (green) were treated with 500 µM IAA for 8 or 20 hours and co-stained with anti-Tgβ-tubulin (red) and Hoechst (blue). Guide panels are a merge of Centrin1 and β-tubulin signals, numbers indicate number of centrosomes. Scale bars: 2 μm. (C-D) Quantification of parasites with single, duplicated, or overduplicated centrosomes (TgCentrin1) grown in the presence or absence of IAA for 8 or 20 hours. Mean±SD of n=3 biological replicates; 200-500 parasites counted per condition.

## Discussion

We have identified TgMAPK2 as essential to progressing through an early checkpoint in the *Toxoplasma* cell cycle. Our data demonstrate that without TgMAPK2, parasites arrest before centrosome duplication, causing a block in the initiation of parasite budding, and eventual parasite death. Intriguingly, organelles, including the Golgi apparatus, apicoplasts, and mitochondria all continue to replicate, and relative parasite size increases, even though budding does not occur (Figure 10A,B). The process of endodyogeny has historically been described as a careful coordination of organellar biogenesis with parasite budding (6, 14, 35). Our data suggest that organelle biogenesis and cell growth are not directly coupled to cell cycle through regulatory checkpoints, but rather through simple timing. This idea is borne out by the ability to produce otherwise normal parasites in which the apicoplast does not replicate (36, 37). In addition, while the residual body is usually first observed after the first round of replication (38), and has recently been linked to inter-parasite trafficking within a vacuole (28), we observed enlarged residual bodies in vacuoles containing single, arrested parasites after long-term (20 h) TgMAPK2 degradation. Our data are therefore consistent with the role of the residual body in response to cellular stress. Notably, treatment of parasites with pyrrolidine dithiocarbamate blocks *Toxoplasma* growth and leads to enlarged residual bodies, which was interpreted as a mechanism of cell size control (39).

**Figure 10.**
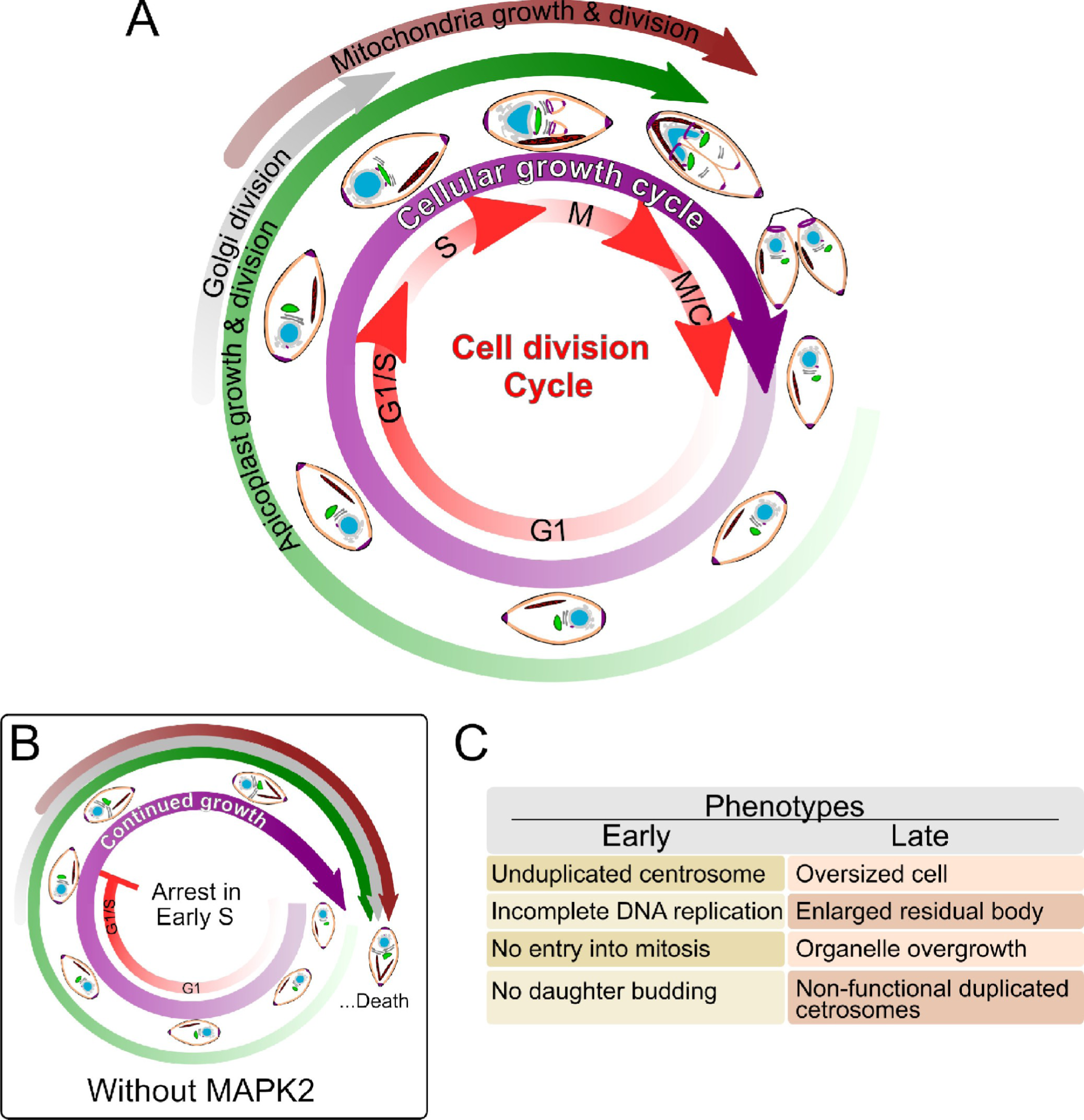
Loss of TgMAPK2 uncouples *Toxoplasma* organelle growth from cell division progression. (A) In normal parasites, cellular and organellar growth is synchronized with the division cycle. Cartoons representations indicate approximate parasite morphologies through the cell cycle. Approximate timing of organellar division are indicated by the colored arrows in the outer circles, and are adapted from Nishi, *et al*. (14). (B) In MAPK2-depleted parasites, parasites appear to arrest in early S phase of the cell cycle, but continue growing and increasing the organelle load. (C) Table summarizing early (8 h) and late (18+ h) appearing phenotypes observed upon MAPK2 depletion.

While depletion of TgMAPK2 results in a reversible arrest just prior to centrosome duplication, after longer periods of growth without TgMAPK2, parasites often had multiple Centrin1-positive puncta (Figure 9), which appear to represent partially formed or otherwise non-functional centrosomes. While these puncta were associated with microtubules, the tubulin structures were not well-organized as in normal parasites (Figure 9). Similarly, we did not observe these puncta associated with the parasite nuclei or organelles. This suggests a partial escape from arrest in which parasites lacking MAPK2 progress down an abnormal and lethal pathway. Similar scenarios have been well documented in cell cycle mutants in other organisms (40, 41).

A number of kinases with orthologs in model organisms have been identified as critical to the *Toxoplasma* cell cycle. These include kinases in the CDK (5, 16), Aurora (17, 42), and NEK (8) families. Progression through G1/S is a critical control point throughout eukaryota, and parasite kinases appear to perform similar functions to their metazoan orthologs in regulating this checkpoint. Notably, TgCRK1, an ortholog of metazoan CDK11, appears to regulate the transcriptional program that allows progress through the G1 checkpoint and centrosome duplication (5). In addition, TgNEK-1 is essential for centrosome separation (8). However, the ortholog of its mammalian substrate, CEP250, does not appear to be a substrate of TgNEK-1. In fact, CEP250 and TgNEK-1 show distinct phenotypes upon disruption (4). Thus, while phylogenetically orthologous proteins to those in well-studied models exist in apicomplexan parasites, they may not be *functionally* orthologous.

In addition to well-conserved kinases, there are a number of Apicomplexa-specific members of many families that typically control the eukaryotic cell cycle (5, 43, 44). In *Toxoplasma*, these include two specialized MAPKs. MAPKL1 is found exclusively in coccidian organisms, which all replicate by endodyogeny during their asexual cycles. MAPK2 is conserved among all extant Alveolata for which genomes are available. While TgMAPKL1 localizes exclusively to the centrosome and prevents its overduplication (3), we have found that TgMAPK2 is required to complete a single round of centrosome duplication. We also found that TgMAPK2 never localizes to the centrosome, nor to the nucleus. It thus seems likely that TgMAPK2 regulates centrosome duplication indirectly, by controlling another process at a distal site that must be completed to progress through this checkpoint. The one unifying feature of all Alveolate organisms is the membrane and cytoskeletal structure known as the inner membrane complex (IMC) in Apicomplexa, and as “alveoli” in other organisms. The processes that regulate and drive the IMC biogenesis are still a mystery. However, the IMC forms the scaffold for new daughter cells (6, 34), a process that is thought to be coordinated by the recently duplicated centrosome (8–10, 12, 34), Given its evolutionary history, it is intriguing to hypothesize that TgMAPK2 plays a role in the crosstalk between these two processes.

## Materials and Methods

### Sequence analysis and phylogeny

Protein sequences for kinases were obtained from ToxoDBv43 and UniProt. The kinase domains were aligned using MAFFT (45), and the resulting alignments were used to estimate the maximum likelihood phylogenetic tree with bootstrap analysis (1000 replicates) in IQ-tree v1.6.12 (46, 47), with the gamma-corrected empirical-frequency LG substitution model.

### PCR and plasmid generation

All PCRs were conducted using Q5 DNA polymerase (New England Biolabs) and the primers listed in Supplemental Table 1. Constructs were assembled using Gibson master mix (New England Biolabs).

### Parasite culture and transfection

Human foreskin fibroblasts (HFFs) were grown in DMEM supplemented with 10% fetal bovine serum and 2 mM glutamine. Toxoplasma tachyzoites were maintained in confluent monolayers of HFF. TgMAPK^3xHA^ and TgMAPK2^AID^ strains were generated by transfecting the RHΔku80Δhxgprt strain (48) or the same strain expressing OsTIR1 driven by the gra1-promoter (22). Transfections included a Cas9/CRISPR vector targeting the TgMAPK2 or TgMAPKL1 3’ and a Q5 PCR product with 500 bp homology arms flanking the appropriate tag together with 10 µg of a Cas9 plasmid. TgMAPK2 complement parasites were created by targeting 3xHA-tagged TgMAPK2 (WT or kinase-dead) driven by DHFR promoter, together with a HXGPRT cassette, to the empty Ku80 locus. mTFP1-α-Tubulin, GFP-α-Tubulin, TgIMC1-mVenus, mVenus-TgCentrin1, GRASP55-mRuby3, or TgEB1-mRuby3 expressing parasites were created by amplifying the FP-marker expression cassette and an adjacent chloramphenicol (or HXGPRT) resistance cassette by PCR and targeting it to a site adjacent Ku80 locus by CRISPR/Cas9-mediated homologous recombination (see Supplemental Table S1) and selecting with chloramphenicol (or MPA/ Xan). TgCEP250L1^3xHA^ parasite was generated by C-terminal single homologous recombination as described (3). The original pTub and pMIN vector was a kind gift of Ke Hu (University of Indiana). The original GRASP55-GFP is driven by pTub promoter, and Neon-Rab5a, EmFP-Rab6, Neon-Rab7, Emerald-Centrin1 and eGFP-Centrin2 are all driven by pMIN promoter, and it was a kind gift of Aoife Heaslip (University of Connecticut).

### Western blotting

Proteins were separated by SDS–PAGE and transferred to a polyvinylidene difluoride membrane. Membranes were blocked for 1 h in PBS + 5% milk, followed by overnight incubation at 4°C with primary antibody in blocking solution. The next day, membranes were washed three times with TBST, followed by incubation at room temperature for 1–2 h with HRP-conjugated secondary antibody (Sigma) in blocking buffer. After three washes with TBST, Western blots were imaged using ECL Plus reagent (Pierce) on a GE ImageQuant LAS4000. Antibodies used in this study include Rb anti-Tgβ-tubulin (1:10000 dilution), rat anti-HA (Sigma; 1:1000 dilution), and mouse m2 anti-FLAG (Sigma; 1:1000 dilution).

### Immunofluorescence and image analysis

HFF cells were grown on coverslips in 24-well plates until confluent and were infected with parasites. The cells were rinsed twice with PBS, and were fixed with 4% paraformaldehyde (PFA)/4% sucrose in PBS at room temperature for 15 min. After two washes with PBS, cells were permeabilized with 0.1% Triton X-100 for 15 min and washed three times with PBS. After blocking in PBS + 3% bovine serum albumin for 30 min, cells were incubated in primary antibody in blocking solution overnight at room temperature. Cells were then washed three times with PBS and incubated with Alexa-Fluor conjugated secondary antibodies (Molecular Probes) for 2 h. Cells were then washed three times with PBS and then mounted with mounting medium containing DAPI (Vector Laboratories). For tyramide amplification of TgMAPK2^AID-3xFLAG^ signal, the above protocol was altered as follows. Endogenous peroxidase activity was quenched by incubation of fixed coverslips with 100 mM sodium azide in PBS for 45 min at room temperature. Cells were blocked with 5% horse serum/0.5% Roche Western Blocking Reagent in TBST for 45 min. HRP-conjugated goat anti-mouse secondary antibody (Sigma) was used and tyramide-fluorophore was allowed to react for 30 s before final washes. Cells were imaged on either a Nikon A1 Laser Scanning Confocal Microscope or a Nikon Ti2E wide-field microscope. Primary antibodies used in this study include rat anti-HA 3F10 (Roche #11867423001), mouse m2 anti-FLAG (Sigma #F1804), rabbit anti-Tgβ-tubulin (1:10,000 dilution), rabbit anti-TOM40 (1:2,000 dilution), rabbit anti-ACP (1:2,000 dilution), rabbit anti-ROP2 (1:10,000 dilution), mouse MAb 45.36 anti-TgIMC1 (1:2,000 dilution, a generous gift from Gary Ward, The University of Vermont). Pearson’s coefficient was calculated for all the Z-stacks of the images of a minimum of 20 cells using Coloc 2 software in ImageJ for each of the markers in Figure 2A,B. Quantitative image analysis in Figure 8 and Supplemental Figure S2 was performed in ImageJ, as follows. All channels were background subtracted and individual parasites were identified manually using IMC1 signal. Total intensity of ACP1, IMC1, and TOM40 signal and cell area were quantified and correlated on a per-cell basis.

### Membrane extraction analysis

Freeze-thaw lysate from TgMAPK2^3xHA^ parasites was ultracentrifuged at 120,000 g for 2 h to separate the soluble supernatant and pellet. The pellets were resuspended and extracted in either PBS; 0.1 M Na_2_CO_3_, pH 11.5; 1 M NaCl; or PBS + 1% Triton X-100 at 4°C for 30 minutes, followed by 120,000 g ultracentrifugation. Samples were analyzed by western blot as described above.

### Plaque assay

Plaque assays were performed using 6-well plates containing HFFs infected with 200 parasites per well in the presence or absence of 500 μM IAA. After 7 days, the cells were fixed with methanol, stained with crystal violet solution, and the resulting plaques were counted. All plaques assays were performed in biological triplicate.

### Transmission electron microscopy

Cells were fixed on MatTek dishes with 2.5% (vol/vol) glutaraldehyde in 0.1 M sodium cacodylate buffer. After three rinses in 0.1 M sodium cacodylate buffer, they were postfixed with 1% osmium tetroxide and 0.8% K_3_[Fe(CN)_6_] in 0.1 M sodium cacodylate buffer for 1 h at room temperature. Cells were rinsed with water and en bloc stained with 2% aqueous uranyl acetate overnight. After three rinses with water, specimens were dehydrated with increasing concentration of ethanol, infiltrated with Embed-812 resin, and polymerized in a 70°C oven overnight. Blocks were sectioned with a diamond knife (Diatome) on a Leica Ultracut UC7 ultramicrotome (Leica Microsystems) and collected onto copper grids, poststained with 2% uranyl acetate in water and lead citrate. All TEM images were acquired on a Tecnai G2 spirit transmission electron microscope (FEI) equipped with a LaB_6_ source at 120 kV. Images in Figure 7B of sectioned cells are representative of 15 TgMAPK2^AID^, 40 TgMAPK2^AID/IAA^ (6h IAA duration) and 40 TgMAPK2^AID/IAA^ (20h IAA duration) vacuoles. The numbers of Golgi per parasite were counted manually from 35 images.

### Invasion and egress assay

Invasion and egress assays were performed using mTFP1-α-Tubulin expressed TgMAPK2^AID^ parasites.) Parasites were allowed to grow overnight without IAA. The next day, 2 h before the assay was initiated, media was switched to IAA (or mock) media. Parasites were mechanically released from host cells, and 2×10^6^ parasites of each condition were added to confluent HFFs grown on coverslips in a 24 well plate, where they were allowed to invade at 37°C for 2 h in -/+ IAA media. These were then washed 10x with PBS, fixed, and attached parasites were stained with anti-SAG1 without permeabilization. Assays were conducted in biological triplicates each of technical triplicates. The mTFP1-α-Tubulin positive but SAG1 negative parasites were regarded as internal (invaded) parasites. To measure egress, parasites were grown for 24–30 h in confluent HFFs on coverslips. 2 h before the assay was initiated, media was switched to IAA, as above. Cells were washed with prewarmed Hank’s balanced salt solution (HBSS) before the assay, and then incubated with HBSS containing 3 μM calcium ionophore A23187 (Cayman Chemical) for 1 min at 37°C before fixation and imaging.

### DNA content analysis by FACS

Parasite nuclear DNA content was determined by FACS using 1 µg/ml DAPI. Intracellular parasites were washed with cold PBS twice, filtered with 5 µm filters, and fixed in ice-cold 80% (vol/vol) ethanol overnight. The parasites were pelleted at 300 g and were resuspended in 1.0 ml freshly-made 1 µg/ml DAPI / Triton X-100 solution at room temperature for 30 min. Data were fit to gaussian mixture models and plotted using the CytoFlow v1.0 python package (49).

### Statistical analysis and figure generation

Statistical tests were conducted in GraphPad Prism v8.4. All shown are mean and error bars indicate S.D. Images were analyzed using the Fiji distribution of ImageJ (50). Data in Figure 8 and Supplement Figure S2 were plotted using the Seaborn python package and statistical tests performed in statsmodels. Figures were created and edited using Inkscape v0.92.

## Acknowledgments

We thank Melanie Cobb and members of the Cobb and Reese labs for lively discussions; the UT Southwestern Electron Microscopy core facility for assistance with data collection. M.L.R. acknowledges funding from the Welch Foundation (I-1936-20170325), National Science Foundation (MCB1553334), and National Institutes of Allergy and Infectious Disease (R01AI150715). X.H. was funded, in part, by Cancer Prevention and Research Institute of Texas Training Grant RP160157. T.B. was funded, in part, by NIH training grant T32GM008203.

---

**Supplemental Figure S1.**
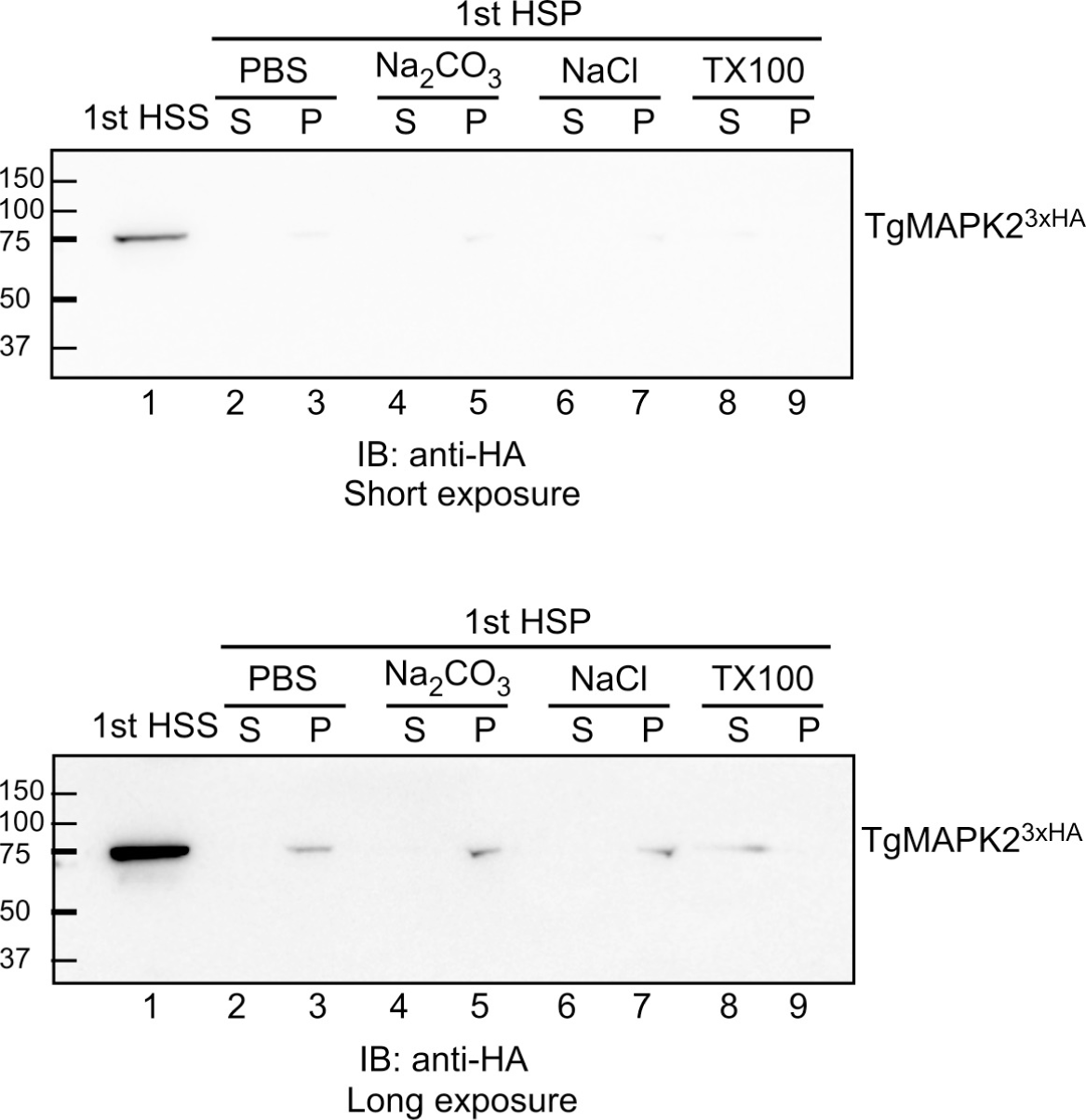
Subcellular fractionation demonstrates that TgMAPK2^3xHA^ behaves as a soluble protein. TgMAPK2^3xHA^ parasites were lysed by freeze-thaw and the lysate was ultracentrifuged at 120,000 g for 2 h to separate the high speed supernatant (1st HSS) and high speed pellet (1st pellet). The pellet was then extracted with either PBS; 0.1 M Na_2_CO_3_, pH 11.5; 1 M NaCl or 1% TritonX-100 (TX100) at 4°C for 30 minutes, resedimented by ultracentrifugation, as above, separated by SDS-PAGE and probed with anti-HA for western blot. Note all lanes represent equivalent relative volumes.

**Supplemental Figure S2.**
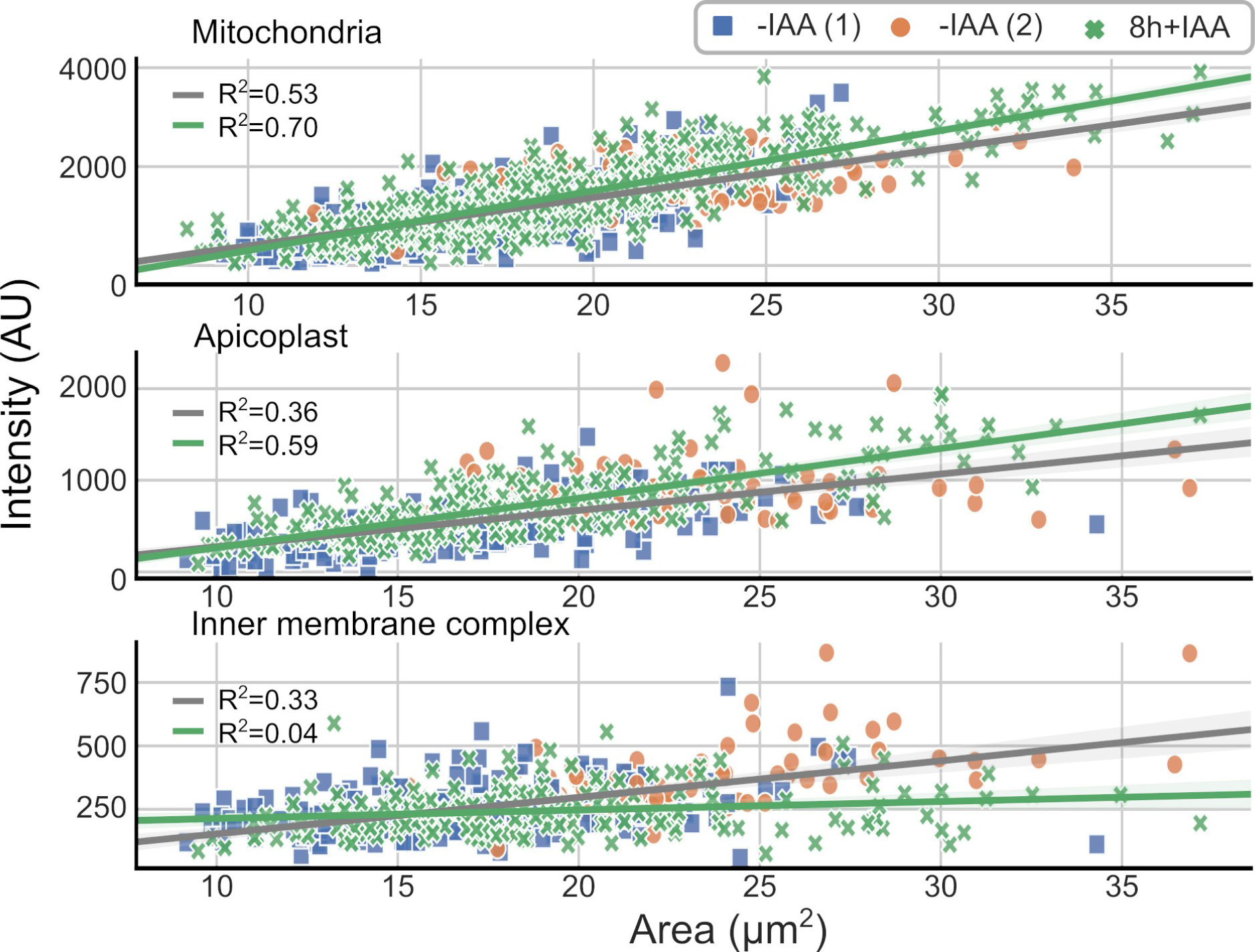
Scatter plot illustrates the total intensity trend of TOM40 (mitochondrion), ACP (apicoplast) and IMC1. -IAA parasites are binned by centrosome number ((1) or (2)), whereas all parasites included in the +IAA dataset had only 1 centrosome. Trendlines represent linear regression for the +IAA (green) and combined -IAA (gray) datasets, shaded bands represents 95% confidence interval. Note the -IAA datasets were combined for linear regression as the one/two centrosome bins tended to cluster in separate quadrants. Data indicate that mitochondrial and apicoplast load, quantified by TOM40 and ACP1 intensity, respectively, was correlated with parasite size. IMC1 intensity, however, correlated more with centrosome numbers than parasite size and was essentially constant in TgMAPK2^AID/IAA^ parasites irrespective of their size (note R^2^=0 for linear regression indicates that a horizontal line is the best model for the data).

